# The DynaSig-ML Python package: automated learning of biomolecular dynamics-function relationships

**DOI:** 10.1101/2022.07.06.499058

**Authors:** Olivier Mailhot, François Major, Rafael Najmanovich

## Abstract

**Summary:** The DynaSig-ML (“Dynamical Signatures - Machine Learning”) Python package allows the efficient, user-friendly exploration of 3D dynamics-function relationships in biomolecules, using datasets of experimental measures from large numbers of sequence variants. The DynaSig-ML package is built around the Elastic Network Contact Model (ENCoM), the first and only sequence-sensitive coarse-grained NMA model, which is used to generate the input Dynamical Signatures. Starting from *in silico* mutated structures, the whole pipeline can be run with just a few lines of Python and modest computational resources. The compute-intensive steps can also easily be parallelized in the case of either large biomolecules or vast amounts of sequence variants. As an example application, we use the DynaSig-ML package to predict the evolutionary fitness of the bacterial enzyme VIM-2 lactamase from deep mutational scan data.

**Availability and implementation:** DynaSig-ML is open source software available at https://github.com/gregorpatof/dynasigml_package

## 1. introduction

The Elastic Network Contact Model (ENCoM) is the only sequence-sensitive coarse-grained normal mode analysis model [1]. Its sequence sensitivity enables its use to predict the impact of sequence variants on biomolecular function through changes in predicted stability [2] and dynamics [3]. We recently extended ENCoM to work on RNA molecules and predicted microRNA maturation efficiency from a dataset of experimentally measured maturation efficiencies of over 26 000 sequence variants [4]. To do so, the ENCoM Dynamical Signatures, which are vectors of predicted structural fluctuations at every position in the system, were used as input variables in a LASSO multiple linear regression model [5] to predict maturation efficiency. To our knowledge, this coupling of coarse-grained normal mode analysis to machine learning in order to predict biomolecular function is the first of its kind. Furthermore, it can be applied to any biomolecule for which there exist experimental data linking perturbations (such as mutations or ligand binding) to function. Indeed, ENCoM is currently applicable to proteins, nucleic acids, small molecules and their complexes [6]. Here we present the DynaSig-ML (“Dynamical Signatures - Machine Learning”) Python package, which allows the implementation and automated replication of that novel protocol. As an example application, we apply DynaSig-ML to predict enzymatic efficiencies of VIM-2 lactamase sequence variants, starting from mutagenesis data. DynaSig-ML automatically computes the ENCoM Dynamical Signatures from a list of perturbed structures (mutations or ligand binding), stores them as lightweight serialized files, and can then be used to train simple machine learning algorithms using the Dynamical Signatures. The first algorithm is LASSO regression, which allows the mapping of the learned coefficients on the studied structure (automatically accomplished by DynaSig-ML plus two simple PyMOL [7] commands). As these coefficients represent the relationship between flexibility change at specific positions and the predicted functional property, this mapping can be used to drive new biological hypotheses. The second machine learning model implemented is the multilayer perceptron (MLP), a type of feedforward neural network [8]. MLPs can learn complex relationships between the input variables and are thus more powerful than LASSO regression, however it is not possible to map the learned patterns back on the structure because of the MLP’s complexity and absence of linear independence between input variables. DynaSig-ML automatically generates graphs showing testing performance. Each of the necessary steps to apply DynaSig-ML is documented online as part of a step by step tutorial (https://dynasigml.readthedocs.io).

## 2. implementation

DynaSig-ML runs the ENCoM model within NRGTEN, another user-friendly, extensively documented Python package [6]. The machine learning models are implemented using the scikit-learn Python package [9]. The numerical computing is accomplished by NumPy [10] and the performance graphs are generated with matplotlib [11], making these four packages the only dependencies of DynaSig-ML.

## 3. vim-2 lactamase example

In order to illustrate a typical use case of DynaSig-ML, we applied it to study dynamics-function relationships from deep mutational scan (DMS) data on the VIM-2 lactamase enzyme [12]. VIM-2 (Verona integron-encoded metallo-*β*-lactamase 2) is a bacterial enzyme capable of degrading *β*-lactam antibiotics, and represents a major source of worldwide antibiotic resistance [13]. The DMS dataset used measured bacterial fitness for every VIM-2 sequence variant under various concentrations of antibiotics [12]. For this application, we use the fitness under the maximal concentration of ampicillin (128*µ*g/mL) at 37 degrees Celsius as the property the machine learning models try to predict from the Dynamical Signatures. Figure 1 illustrates the whole protocol used to start from the PDB [14] structure of VIM-2 lactamase [15], train the machine learning models, test their performance and map the LASSO coefficients back on the VIM-2 structure.

**Figure 1:**
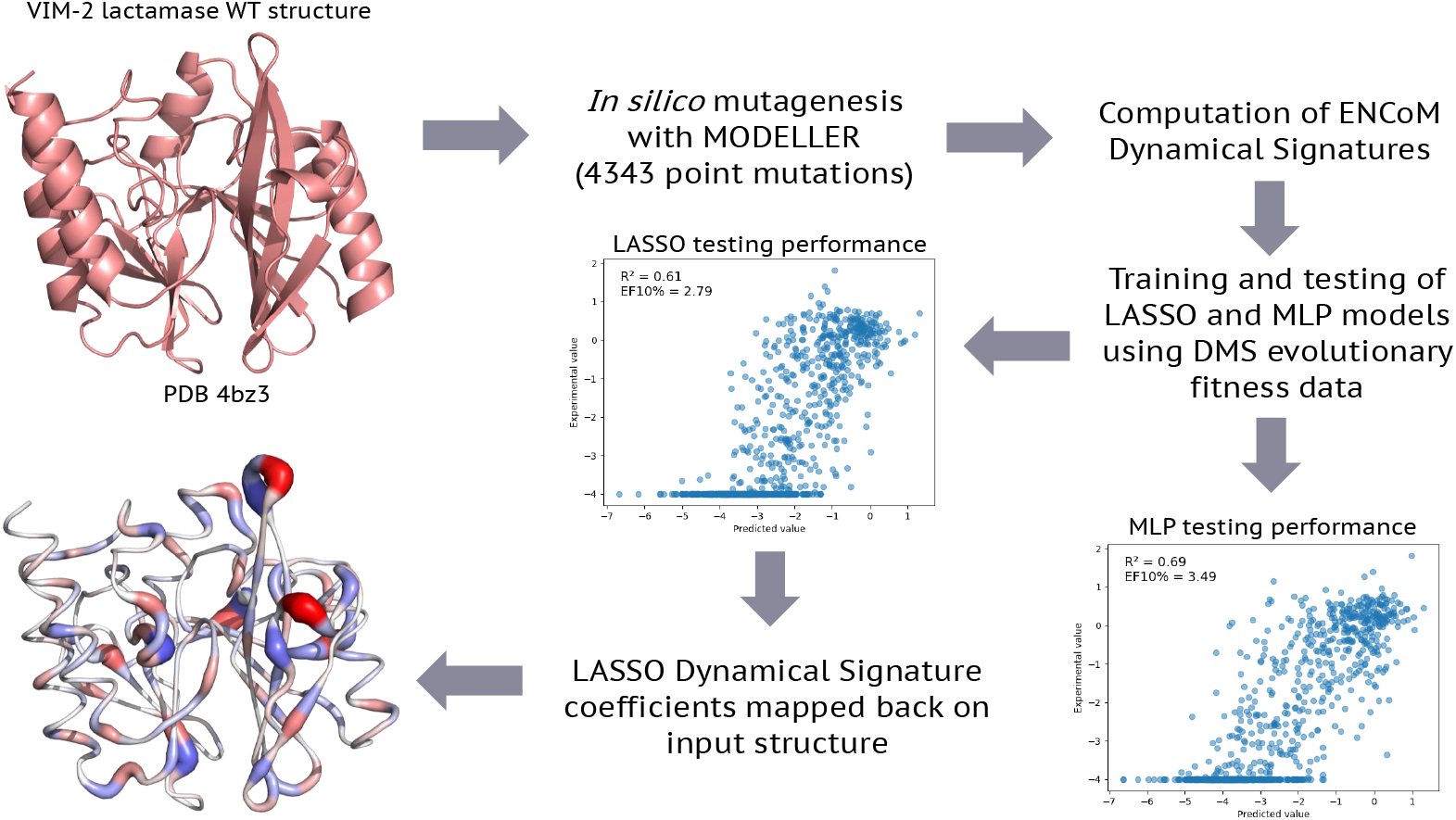
ENCoM-DynaSig-ML pipeline applied to VIM-2 lactamase deep mutational scan data. The crystal structure of VIM-2 lactamase is used as a template to perform the 4343 point mutations with experimental fitness data using the MODELLER software [16], all subsequent steps are performed using DynaSig-ML. For each of the in silico variants, a Dynamical Signature is computed with ENCoM. LASSO regression and multilayer perceptron models are trained using as input variables the Dynamical Signatures and the structural attributes used by Chen et al. as part of the VIM-2 deep mutational scan study [12]. In the case of the LASSO regression model, the independence of the input variables allows the mapping of the learned coefficients back on the VIM-2 structure. The color gradient represents each coefficient, from blue for negative coefficients, to white for null coefficients and red for positive coefficients. The largest absolute value coefficient will have the brightest color. The sign of a coefficient captures the nature of the relationship between flexibility changes at that position and the experimental property of interest (in this case, evolutionary fitness). Negative coefficients mean that rigidification of the position leads to higher fitness, while positive coefficients mean that softening of that position leads to higher fitness. The thickness of the cartoon represents the absolute value of the coefficients, i.e. their relative importance in the model.

Interestingly, the Dynamical Signatures exhibit good complementarity to the other attributes used by Chen *et al*., which are the mean ΔΔG of folding calculated with Rosetta [17], the change in accessible surface area, and a vector of length 40 which specifies what are the starting and mutated amino acid identities for the point mutation. The authors report a training coefficient of determination of R^2^ = 0.55 when fitting a linear model with these input variables to all the available data. They do not report testing performance, so this 0.55 R^2^ can be seen as an upper bound for the performance of a linear model based on these properties. Since the dataset contains point mutations only, there can be no sequence redundancy between the training and testing set. For this reason, we generated a random 80/20 train/test split for this application. The exact variants randomly picked for the testing set are available in the GitHub repository that accompanies the online DynaSig-ML tutorial (https://github.com/gregorpatof/dynasigml_vim2_example). When combining the Chen *et al*. attributes and Dynamical Signatures, we obtain LASSO and MLP models reaching respective testing performances of R^2^ = 0.61 and R^2^ = 0.69. The enrichment factors at 10%, which are values ranging from 0 to 10 characterizing the relative proportion of the top 10% measured values in the top 10% predicted values, are 2.79 and 3.49 for the LASSO and MLP models respectively. For a more in-depth analysis of performance including models trained with the Dynamical Signatures alone and static predictors alone, see Supplementary Information.

## 4. conclusions

In conclusion, the DynaSig-ML Python package allows the fast and user-friendly exploration of dynamics-function relationships in biomolecules. It uses the ENCoM model, the first and only sequence-sensitive coarse-grained normal mode analysis model, to automatically compute Dynamical Signatures from structures in PDB format, stores them as lightweight serialized Python objects, and automatically trains and tests LASSO regression and MLP models to predict experimental measures. Moreover, it automatically generates performance graphs and maps the LASSO coefficients back on the input PDB structure. A detailed online tutorial is available to replicate the VIM-2 deep mutational scan application presented here (https://dynasigml.readthedocs.io).

## Supporting information

Supplementary data, images and tables

## 5. acknowledgements and funding

RJN is part of PROTEO (the Québec network for research on protein function, structure and engineering).

This work was supported by Natural Sciences and Engineering Research Council of Canada (NSERC) Discovery program grants (FM and RN); Genome Canada and Genome Quebec (RN); Compute Canada (RN) and Canadian Institutes of Health Research (CIHR) (FM, grant number MOP-93679). OM is the recipient of a Fonds de Recherche du Québec—Nature et Technologies (FRQ-NT) Doctorate’s scholarship; and a Faculté des Études Supérieures et Postdoctorales de l’Université de Montréal scholarship for direct passage to the PhD.

