## Supplementary data, images and tables for "The DynaSig-ML Python package: automated learning of biomolecular dynamics-function relationships"

### Supplementary information for: The DynaSig-ML Python package: automated learning of biomolecular dynamics-function relationships


Olivier Mailhot[1−4] François Major[2,3] Rafael Najmanovich[4,*]

[1]Department of Biochemistry and Molecular Medicine, Université de Montréal, Montreal, Canada
[2]Department of Computer Science and Operations Research, Université de Montréal, Montreal, Canada
[3]Institute for Research in Immunology and Cancer, Université de Montréal, Montreal, Canada
[4]Department of Pharmacology and Physiology, Université de Montréal, Montreal, Canada
[*]To whom correspondence should be addressed.


July 6, 2022


Contact:


#### 1 SUPPLEMENTARY INFORMATION

Supplementary Figure 1 reports the testing performance of LASSO and MLP models when trained using the ENCoM Dynamical Signatures combined to the static predictors from Chen *et al.* [1], which are the Rosetta ΔΔG of folding, the change in solvent accessible surface area and 40 variables describing the starting and ending amino acid for the mutation (20 possibilities for each). In order to investigate the contributions of Dynamical Signatures and static descriptors alone, we also trained LASSO and MLP models using either one of these sets of variables alone. The testing performances are shown in Supplementary Figure 2 and Supplementary Figure 3. Interestingly, while the testing $R^2$ is lower for the DynaSig models (0.24 vs 0.52 for LASSO, 0.46 vs 0.58 for MLP), both EF10% are identical (2.79 for LASSO and 3.26 for MLP). This led us to hypothesize that the models might be enriching the same variants as their top predictions, however it is not the case. Supplementary Table 2 lists all variants that in the top 10% for any of the tested models or for the experimentally measured fitness. Surprisingly, despite identical EF10% performances, the DynaSig and static models have few top predictions in common (14 out of 87 for the LASSO models, 29 out of 87 for the MLP models). This discrepancy in ranking the variants might explain the good complementarity of the two sets of features, especially when it comes to predictive $R^2$. Another striking finding is that the DynaSig models seem to benefit more from the power of the MLP, as $R^2$ almost doubles going from LASSO to MLP. This big gain in performance might be explained by the intrisincally nonlinear relationships between flexibility changes at different positions (the whole protein is a coupled system), which cannot be captured by linear regression.

When it comes to predicting experimental fitness for variants containing more than one mutation, the starting and ending amino acid vector used by Chen *et al.* cannot be used. However, from a biomolecular engineering point of view, variants containing multiple mutations are where computational methods really shine as the number of possible variants grows exponentially with the number of mutations. The Dynamical Signatures however can be computed for any sequence variant as long as the assumption that the equilibrium structure does not change considerably holds. We have recently predicted maturation efficiency for miR-125a sequence variants containing up to 6 mutations [2]. In order to investigate what performance one could expect using only the static properties generalizable to variants with multiple mutations, we trained LASSO and MLP models using only the accessible solvent area change and predicted ΔΔG of folding. The testing performances are reported in Supplementary Figure 4 and while the $R^2$ coefficients are fair (0.42 and 0.45 for LASSO and MLP), the EF10% values are lower than with all other predictors used at 1.86 for LASSO and 2.67 for MLP. This illustrates the advantage of the Dynamical Signatures, as they are generalizable to multiple mutations and perform on par with the full static descriptors when it comes to EF10%.

Both training and testing $R^2$ and EF10% are provided in Supplementary Table 1 as a means of quick comparison between the different models. We obtain similar training $R^2$ as Chen *et al.* using their static descriptors (0.54 for our implementation, 0.55 is what was reported), and the combination of Dynamical Signatures and static descriptors reaches training $R^2$ of 0.64 for LASSO and 0.96 for MLP.

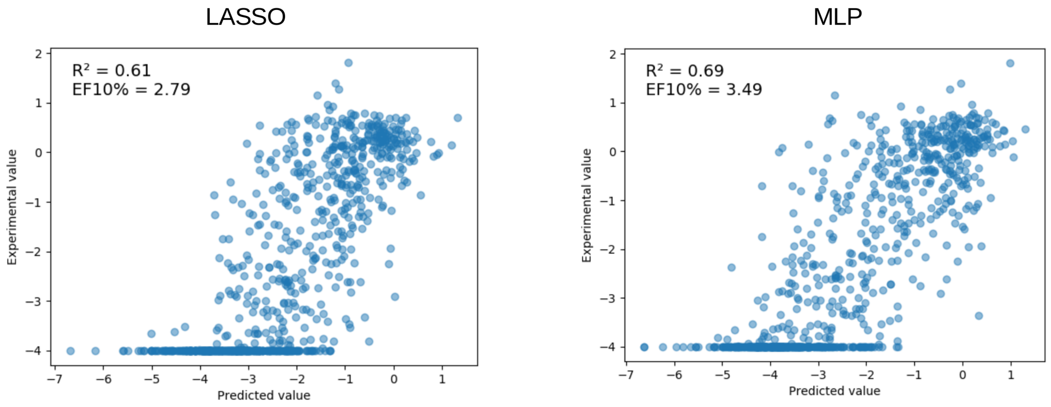


**Supplementary Figure 1: Testing performance using all static descriptors and Dynamical Signatures.**

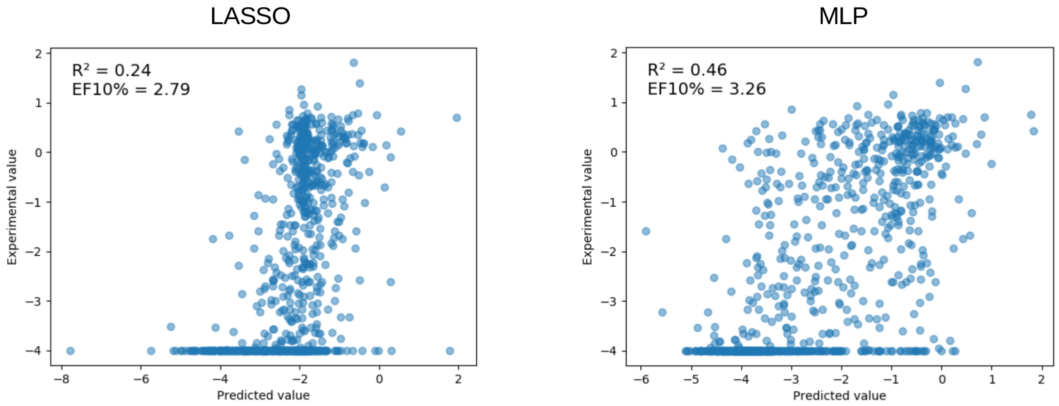


**Supplementary Figure 2: Testing performance using only Dynamical Signatures.**

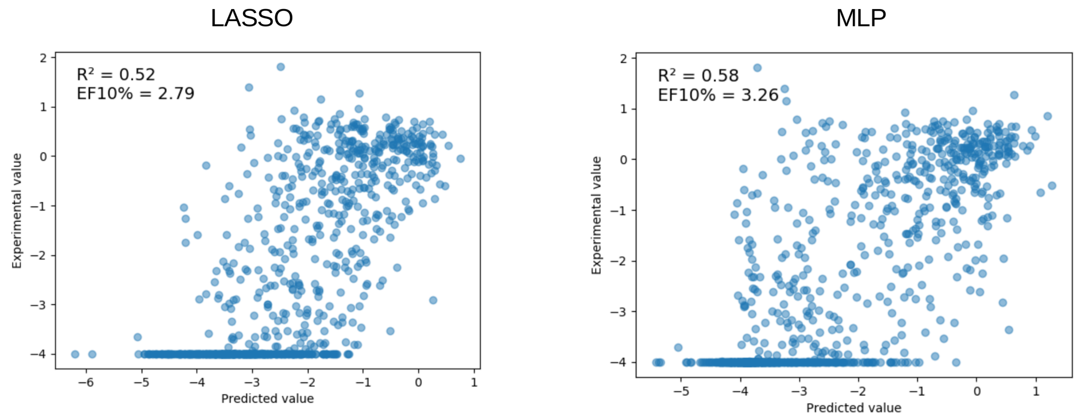


**Supplementary Figure 3: Testing performance using only static descriptors.**

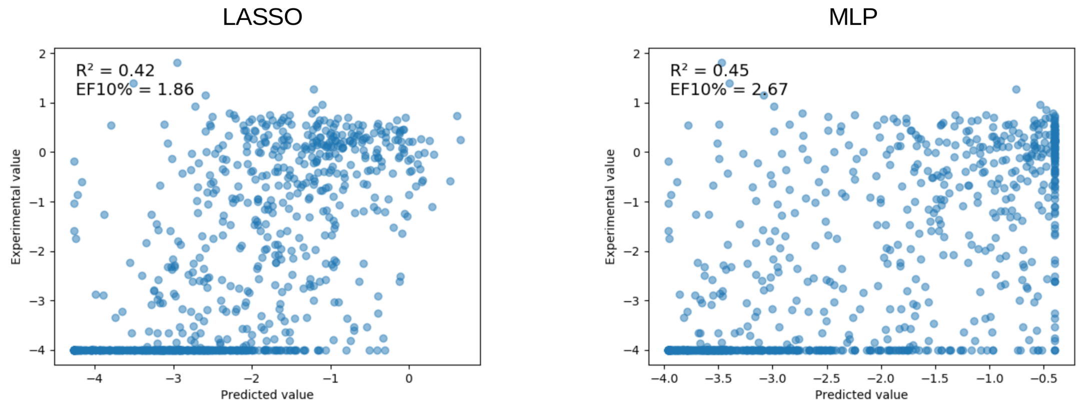


**Supplementary Figure 4: Testing performance using only the solvent accessible area change and folding ΔΔG.**

**Supplementary Table 1:** Testing and training $R^2$ are reported for LASSO and MLP models trained on the combination of Dynamical Signatures and all static descriptors (All), the Dynamical Signatures only (DynaSig), the static descriptors only (Static) and for the combination of folding ΔΔG and accessible solvent area (ASA ΔΔG).

| Training variables | ML model | Testing $R^2$ | Training $R^2$ | EF10% |
|---|---|---|---|---|
| All | LASSO | 0.61 | 0.64 | 2.79 |
| All | MLP | 0.69 | 0.96 | 3.49 |
| DynaSig | LASSO | 0.24 | 0.29 | 2.79 |
| DynaSig | MLP | 0.46 | 0.82 | 3.26 |
| Static | LASSO | 0.52 | 0.54 | 2.79 |
| Static | MLP | 0.58 | 0.78 | 3.26 |
| ASA ddG | LASSO | 0.42 | 0.45 | 1.86 |
| ASA ddG | MLP | 0.45 | 0.50 | 2.67 |

**Supplementary Table 2:** Top 10% testing set predicted single mutants across all tested models. LASSO and MLP top 10% predictions are reported for the combination of Dynamical Signatures and all static descriptors (all), for the Dynamical Signatures only (DynaSig), for the static descriptors only (static) and for the combination of folding ΔΔG and accessible solvent area (ΔΔG ASA). The true top 10% experimental measurements for the testing set are identified in the second column.

| Variant ID | Exp top 10% | LASSO all | MLP all | LASSO DynaSig | MLP DynaSig | LASSO static | MLP static | LASSO ASA ΔΔG | MLP ASA ΔΔG |
|---|---|---|---|---|---|---|---|---|---|
| E32A | FALSE | IN | IN | OUT | OUT | IN | OUT | IN | IN |
| E32I | FALSE | IN | OUT | OUT | OUT | IN | OUT | IN | IN |
| E32M | FALSE | IN | OUT | OUT | OUT | IN | OUT | IN | IN |
| P34F | FALSE | OUT | OUT | IN | OUT | OUT | OUT | OUT | OUT |
| T35E | TRUE | OUT | OUT | IN | IN | OUT | OUT | OUT | OUT |
| T35I | TRUE | OUT | IN | OUT | IN | OUT | OUT | OUT | OUT |
| V36D | TRUE | OUT | OUT | IN | OUT | OUT | OUT | OUT | OUT |
| V36G | TRUE | IN | IN | IN | IN | OUT | OUT | OUT | OUT |
| V36H | TRUE | OUT | OUT | OUT | IN | OUT | OUT | OUT | OUT |
| S37D | FALSE | IN | IN | OUT | OUT | IN | OUT | IN | IN |
| S37E | FALSE | IN | OUT | OUT | OUT | IN | OUT | IN | OUT |
| S37M | FALSE | IN | OUT | OUT | OUT | IN | OUT | IN | IN |
| S37Q | FALSE | IN | OUT | OUT | OUT | IN | OUT | IN | IN |
| E38C | FALSE | OUT | OUT | OUT | IN | OUT | OUT | OUT | IN |
| E38H | FALSE | OUT | IN | OUT | IN | OUT | IN | OUT | OUT |
| E38R | FALSE | OUT | IN | OUT | IN | OUT | IN | OUT | OUT |
| V41G | TRUE | IN | OUT | IN | IN | OUT | OUT | OUT | OUT |
| V41I | FALSE | OUT | OUT | OUT | OUT | OUT | IN | OUT | IN |
| V41K | FALSE | OUT | OUT | OUT | OUT | OUT | IN | OUT | OUT |
| V41L | FALSE | OUT | OUT | OUT | OUT | OUT | IN | OUT | OUT |
| V41M | FALSE | OUT | IN | OUT | OUT | OUT | IN | OUT | OUT |
| V41P | TRUE | OUT | OUT | OUT | OUT | OUT | IN | OUT | OUT |
| V41R | FALSE | IN | OUT | OUT | OUT | OUT | IN | OUT | OUT |
| E43A | FALSE | OUT | IN | OUT | OUT | OUT | IN | OUT | OUT |
| E43H | FALSE | OUT | OUT | OUT | OUT | OUT | IN | OUT | OUT |
| E43I | FALSE | OUT | OUT | OUT | OUT | OUT | OUT | OUT | IN |
| E43K | FALSE | OUT | IN | IN | OUT | OUT | OUT | OUT | IN |
| R45E | FALSE | OUT | OUT | IN | OUT | OUT | OUT | OUT | OUT |
| R45M | TRUE | OUT | OUT | IN | OUT | OUT | OUT | OUT | OUT |
| Y47A | TRUE | OUT | OUT | IN | OUT | OUT | OUT | OUT | OUT |
| Y47G | FALSE | OUT | OUT | IN | OUT | OUT | OUT | OUT | OUT |
| Q48G | FALSE | OUT | OUT | OUT | IN | OUT | OUT | OUT | OUT |
| Q48H | FALSE | OUT | OUT | OUT | OUT | IN | OUT | OUT | IN |
| Q48L | FALSE | IN | OUT | OUT | IN | IN | OUT | OUT | IN |
| Q48P | TRUE | IN | OUT | OUT | OUT | IN | OUT | IN | IN |
| Q48R | TRUE | IN | OUT | IN | OUT | IN | IN | OUT | IN |
| A50K | FALSE | OUT | OUT | OUT | OUT | OUT | IN | OUT | OUT |
| A50Q | FALSE | OUT | OUT | OUT | OUT | OUT | IN | OUT | OUT |
| A50S | FALSE | OUT | IN | OUT | OUT | OUT | IN | OUT | OUT |
| D51E | FALSE | OUT | OUT | OUT | OUT | OUT | OUT | IN | IN |
| D51I | FALSE | OUT | OUT | OUT | OUT | OUT | OUT | IN | OUT |
| D51L | FALSE | OUT | OUT | OUT | OUT | OUT | OUT | IN | IN |
| D51R | FALSE | OUT | OUT | OUT | OUT | OUT | OUT | IN | OUT |
| D51V | FALSE | OUT | OUT | OUT | OUT | IN | OUT | IN | IN |

| Supplementary Table 4 (continued) | | | | | | | | | |
|---|---|---|---|---|---|---|---|---|---|
| Variant ID | Exp top 10% | LASSO all | MLP all | LASSO DynaSig | MLP DynaSig | LASSO static | MLP static | LASSO ASA ΔΔG | MLP ASA ΔΔG |
| S55A | TRUE | OUT | IN | IN | IN | OUT | OUT | OUT | OUT |
| S55F | FALSE | OUT | OUT | IN | OUT | OUT | OUT | OUT | OUT |
| S55M | TRUE | OUT | IN | IN | IN | OUT | OUT | OUT | OUT |
| S55P | FALSE | OUT | OUT | IN | OUT | OUT | OUT | OUT | OUT |
| S61F | FALSE | OUT | OUT | OUT | OUT | OUT | OUT | OUT | IN |
| S61N | FALSE | IN | OUT | OUT | OUT | IN | IN | IN | OUT |
| D63G | TRUE | OUT | OUT | OUT | OUT | IN | OUT | IN | OUT |
| D63I | FALSE | OUT | OUT | OUT | OUT | OUT | OUT | IN | OUT |
| D63L | FALSE | OUT | OUT | OUT | OUT | IN | OUT | IN | IN |
| D63N | FALSE | IN | OUT | OUT | OUT | IN | IN | IN | OUT |
| D63W | FALSE | OUT | OUT | OUT | OUT | OUT | OUT | IN | IN |
| G64E | FALSE | OUT | OUT | OUT | OUT | OUT | OUT | IN | OUT |
| A65G | FALSE | OUT | IN | OUT | OUT | OUT | OUT | OUT | OUT |
| A65H | TRUE | OUT | IN | OUT | OUT | OUT | OUT | OUT | OUT |
| A65Y | TRUE | OUT | IN | OUT | OUT | OUT | IN | OUT | OUT |
| V66F | TRUE | OUT | IN | OUT | OUT | OUT | OUT | OUT | OUT |
| V66R | TRUE | OUT | OUT | OUT | IN | OUT | IN | OUT | OUT |
| V66Y | TRUE | OUT | OUT | OUT | OUT | OUT | IN | OUT | OUT |
| Y67A | TRUE | OUT | OUT | IN | OUT | OUT | OUT | OUT | OUT |
| Y67H | FALSE | OUT | IN | OUT | OUT | OUT | IN | OUT | OUT |
| P68C | FALSE | OUT | OUT | IN | OUT | OUT | OUT | OUT | OUT |
| P68Y | TRUE | OUT | OUT | IN | IN | OUT | OUT | OUT | OUT |
| D78C | FALSE | OUT | OUT | OUT | OUT | OUT | OUT | IN | OUT |
| D78M | FALSE | OUT | OUT | OUT | OUT | OUT | OUT | IN | OUT |
| D78P | FALSE | OUT | OUT | OUT | OUT | OUT | OUT | IN | OUT |
| D78Q | FALSE | OUT | OUT | OUT | OUT | OUT | OUT | IN | OUT |
| D78V | FALSE | OUT | OUT | OUT | OUT | OUT | OUT | IN | OUT |
| D78W | FALSE | OUT | OUT | OUT | IN | OUT | OUT | IN | OUT |
| E79C | FALSE | OUT | OUT | IN | OUT | OUT | OUT | OUT | OUT |
| E79G | FALSE | OUT | OUT | IN | OUT | OUT | OUT | OUT | IN |
| I83F | FALSE | OUT | OUT | IN | OUT | OUT | OUT | OUT | OUT |
| A89H | FALSE | OUT | IN | OUT | OUT | OUT | OUT | OUT | OUT |
| K90A | TRUE | IN | IN | OUT | IN | IN | IN | OUT | OUT |
| A93L | TRUE | OUT | IN | OUT | IN | IN | OUT | OUT | OUT |
| A93T | TRUE | OUT | IN | OUT | OUT | OUT | IN | OUT | OUT |
| A94E | FALSE | OUT | IN | OUT | OUT | OUT | OUT | OUT | OUT |
| A97I | FALSE | IN | OUT | OUT | OUT | IN | OUT | OUT | OUT |
| A97Q | FALSE | IN | IN | OUT | OUT | IN | IN | OUT | IN |
| E98Y | FALSE | OUT | OUT | IN | OUT | OUT | OUT | OUT | OUT |
| I99W | FALSE | OUT | OUT | IN | OUT | OUT | OUT | OUT | OUT |
| E100M | FALSE | IN | IN | OUT | OUT | IN | IN | IN | OUT |
| E100R | TRUE | OUT | IN | OUT | OUT | OUT | IN | OUT | OUT |
| E100W | FALSE | OUT | IN | OUT | OUT | OUT | IN | IN | IN |
| K101F | FALSE | IN | OUT | OUT | IN | IN | OUT | IN | OUT |
| K101Y | FALSE | IN | OUT | OUT | IN | IN | OUT | IN | OUT |
| Q102H | TRUE | IN | OUT | IN | OUT | OUT | OUT | OUT | OUT |
| Q102K | TRUE | OUT | IN | OUT | IN | OUT | IN | OUT | OUT |
| Q102L | TRUE | IN | IN | OUT | IN | IN | OUT | OUT | IN |

| Supplementary Table 4 (continued) | | | | | | | | | |
|---|---|---|---|---|---|---|---|---|---|
| Variant ID | Exp top 10% | LASSO all | MLP all | LASSO DynaSig | MLP DynaSig | LASSO static | MLP static | LASSO ASA ΔΔG | MLP ASA ΔΔG |
| Q102T | TRUE | IN | IN | OUT | OUT | OUT | IN | OUT | OUT |
| L105A | FALSE | OUT | OUT | IN | OUT | OUT | OUT | OUT | OUT |
| L105S | FALSE | OUT | OUT | IN | OUT | OUT | OUT | OUT | OUT |
| R109A | FALSE | OUT | IN | OUT | OUT | OUT | OUT | OUT | OUT |
| T113M | FALSE | OUT | OUT | IN | OUT | OUT | OUT | OUT | OUT |
| H114E | FALSE | OUT | OUT | OUT | OUT | OUT | OUT | IN | OUT |
| F115Y | FALSE | OUT | OUT | IN | OUT | OUT | OUT | OUT | OUT |
| D117L | FALSE | OUT | OUT | OUT | OUT | OUT | OUT | OUT | IN |
| D117N | FALSE | OUT | OUT | OUT | OUT | OUT | OUT | OUT | IN |
| D117V | FALSE | OUT | OUT | OUT | OUT | OUT | OUT | OUT | IN |
| G121D | FALSE | OUT | IN | OUT | OUT | OUT | OUT | OUT | OUT |
| D124F | FALSE | OUT | OUT | OUT | OUT | OUT | OUT | OUT | IN |
| D124N | FALSE | OUT | IN | OUT | OUT | OUT | OUT | IN | IN |
| V125M | FALSE | OUT | IN | OUT | OUT | OUT | IN | OUT | OUT |
| R127E | FALSE | OUT | OUT | IN | IN | OUT | OUT | OUT | OUT |
| R127T | FALSE | OUT | OUT | OUT | IN | OUT | OUT | OUT | OUT |
| A128C | FALSE | OUT | OUT | OUT | OUT | OUT | OUT | IN | OUT |
| A128R | FALSE | IN | OUT | OUT | OUT | IN | OUT | IN | IN |
| A129C | FALSE | OUT | OUT | OUT | IN | OUT | OUT | OUT | OUT |
| A129E | FALSE | OUT | OUT | OUT | IN | OUT | IN | OUT | OUT |
| A129G | FALSE | IN | IN | OUT | OUT | OUT | IN | OUT | OUT |
| A129S | FALSE | IN | IN | OUT | OUT | OUT | IN | OUT | OUT |
| A129V | FALSE | OUT | OUT | OUT | IN | OUT | OUT | OUT | OUT |
| A132K | TRUE | OUT | IN | OUT | OUT | OUT | IN | OUT | OUT |
| A132M | TRUE | OUT | IN | OUT | OUT | OUT | OUT | OUT | OUT |
| A132N | FALSE | IN | IN | OUT | OUT | IN | IN | OUT | OUT |
| P137K | FALSE | OUT | OUT | OUT | IN | OUT | OUT | OUT | OUT |
| S138C | FALSE | OUT | OUT | OUT | OUT | OUT | OUT | OUT | IN |
| S138K | FALSE | OUT | OUT | IN | OUT | OUT | OUT | OUT | OUT |
| S138L | FALSE | OUT | OUT | OUT | OUT | OUT | OUT | OUT | IN |
| S138T | FALSE | IN | OUT | OUT | OUT | OUT | OUT | OUT | IN |
| S138Y | FALSE | OUT | OUT | IN | OUT | OUT | OUT | OUT | OUT |
| R141H | TRUE | OUT | OUT | OUT | OUT | IN | OUT | IN | IN |
| R141K | TRUE | IN | OUT | OUT | OUT | IN | IN | IN | OUT |
| R141M | FALSE | IN | IN | OUT | OUT | IN | IN | IN | IN |
| E144K | TRUE | IN | IN | OUT | OUT | IN | IN | IN | IN |
| E144V | FALSE | OUT | OUT | OUT | OUT | IN | OUT | OUT | OUT |
| V145D | TRUE | OUT | IN | OUT | IN | OUT | IN | IN | IN |
| V145K | TRUE | IN | OUT | OUT | IN | IN | IN | IN | IN |
| E146A | TRUE | OUT | IN | OUT | IN | OUT | IN | IN | IN |
| E146I | FALSE | OUT | OUT | IN | OUT | OUT | OUT | OUT | OUT |
| E146R | FALSE | IN | IN | OUT | IN | OUT | IN | OUT | OUT |
| G147N | FALSE | OUT | OUT | OUT | OUT | OUT | OUT | IN | OUT |
| N148E | FALSE | OUT | OUT | OUT | IN | OUT | OUT | OUT | OUT |
| N148M | TRUE | OUT | IN | OUT | OUT | OUT | OUT | OUT | OUT |
| E149R | FALSE | OUT | OUT | IN | OUT | OUT | OUT | OUT | OUT |
| I150V | FALSE | OUT | OUT | OUT | OUT | OUT | IN | OUT | OUT |
| H153P | FALSE | OUT | OUT | IN | OUT | OUT | OUT | OUT | OUT |

| Supplementary Table 4 (continued) | | | | | | | | | |
|---|---|---|---|---|---|---|---|---|---|
| Variant ID | Exp top 10% | LASSO all | MLP all | LASSO DynaSig | MLP DynaSig | LASSO static | MLP static | LASSO ASA ΔΔG | MLP ASA ΔΔG |
| H153S | FALSE | OUT | OUT | IN | OUT | OUT | OUT | OUT | OUT |
| S154A | TRUE | OUT | IN | OUT | OUT | OUT | IN | OUT | OUT |
| S154D | FALSE | OUT | OUT | OUT | OUT | OUT | OUT | OUT | IN |
| E156I | FALSE | OUT | OUT | OUT | IN | IN | OUT | IN | IN |
| E156P | FALSE | IN | OUT | OUT | IN | IN | OUT | IN | IN |
| E156S | FALSE | IN | OUT | OUT | IN | IN | IN | IN | IN |
| S159D | TRUE | IN | OUT | IN | OUT | IN | OUT | IN | IN |
| S159E | TRUE | IN | OUT | IN | OUT | IN | OUT | IN | IN |
| S159N | FALSE | IN | IN | OUT | OUT | IN | OUT | IN | IN |
| S160D | FALSE | OUT | IN | OUT | OUT | OUT | IN | IN | IN |
| S160K | TRUE | IN | OUT | OUT | IN | IN | OUT | IN | IN |
| S161W | FALSE | OUT | IN | OUT | IN | OUT | OUT | OUT | OUT |
| A164D | FALSE | OUT | OUT | OUT | IN | OUT | OUT | OUT | OUT |
| A164I | FALSE | OUT | OUT | OUT | IN | OUT | OUT | OUT | OUT |
| A164L | TRUE | OUT | OUT | OUT | IN | OUT | OUT | OUT | OUT |
| A164M | TRUE | OUT | IN | IN | IN | OUT | OUT | OUT | OUT |
| A164Q | FALSE | IN | IN | OUT | IN | OUT | IN | OUT | OUT |
| V165I | FALSE | OUT | OUT | OUT | OUT | OUT | IN | OUT | OUT |
| V165M | FALSE | OUT | OUT | OUT | OUT | OUT | IN | OUT | OUT |
| R166Q | TRUE | IN | IN | IN | IN | OUT | IN | OUT | IN |
| R166V | FALSE | IN | IN | IN | IN | IN | OUT | IN | IN |
| V170F | FALSE | OUT | OUT | IN | OUT | OUT | OUT | OUT | OUT |
| V170M | FALSE | OUT | OUT | IN | OUT | OUT | OUT | OUT | OUT |
| A177L | FALSE | OUT | OUT | OUT | IN | OUT | OUT | OUT | OUT |
| A177Q | FALSE | OUT | OUT | IN | OUT | OUT | OUT | OUT | OUT |
| A177Y | FALSE | OUT | OUT | OUT | IN | OUT | OUT | OUT | OUT |
| A178G | FALSE | OUT | OUT | IN | IN | OUT | OUT | OUT | OUT |
| T181D | FALSE | IN | OUT | IN | OUT | OUT | IN | OUT | OUT |
| T181E | FALSE | OUT | IN | OUT | IN | OUT | IN | OUT | IN |
| T181H | FALSE | OUT | IN | OUT | IN | OUT | IN | OUT | OUT |
| T181L | TRUE | OUT | OUT | OUT | OUT | IN | OUT | OUT | IN |
| T181R | FALSE | IN | IN | OUT | IN | IN | OUT | OUT | IN |
| T181V | FALSE | IN | OUT | OUT | OUT | IN | IN | OUT | OUT |
| T181Y | FALSE | IN | OUT | IN | OUT | OUT | OUT | OUT | OUT |
| L184M | FALSE | OUT | OUT | IN | OUT | OUT | OUT | OUT | OUT |
| I185V | TRUE | OUT | OUT | OUT | OUT | OUT | IN | OUT | OUT |
| S190D | FALSE | IN | OUT | OUT | OUT | IN | OUT | IN | IN |
| S190E | FALSE | IN | OUT | OUT | OUT | IN | OUT | IN | IN |
| S190H | TRUE | IN | OUT | OUT | IN | IN | OUT | IN | OUT |
| S190K | FALSE | IN | OUT | OUT | OUT | IN | OUT | IN | OUT |
| S190N | FALSE | IN | OUT | OUT | OUT | IN | OUT | IN | IN |
| S190Y | FALSE | IN | OUT | OUT | OUT | IN | OUT | IN | IN |
| S192A | TRUE | OUT | IN | OUT | IN | OUT | IN | OUT | OUT |
| S192H | FALSE | OUT | IN | OUT | OUT | OUT | OUT | OUT | OUT |
| S192N | FALSE | OUT | OUT | OUT | OUT | OUT | IN | OUT | OUT |
| S192T | FALSE | OUT | OUT | OUT | OUT | OUT | IN | OUT | OUT |
| V193K | FALSE | OUT | OUT | OUT | IN | OUT | OUT | OUT | OUT |
| V193R | FALSE | OUT | OUT | OUT | IN | OUT | OUT | OUT | OUT |

| Supplementary Table 4 (continued) | | | | | | | | | |
|---|---|---|---|---|---|---|---|---|---|
| Variant ID | Exp top 10% | LASSO all | MLP all | LASSO DynaSig | MLP DynaSig | LASSO static | MLP static | LASSO ASA ΔΔG | MLP ASA ΔΔG |
| A199C | FALSE | OUT | OUT | IN | OUT | OUT | OUT | OUT | OUT |
| A199F | FALSE | OUT | OUT | IN | OUT | OUT | OUT | OUT | OUT |
| I200V | TRUE | OUT | IN | OUT | OUT | OUT | IN | OUT | OUT |
| L203F | FALSE | OUT | OUT | IN | OUT | OUT | OUT | OUT | OUT |
| L203G | FALSE | OUT | OUT | IN | IN | OUT | OUT | OUT | OUT |
| S204D | FALSE | IN | IN | OUT | OUT | IN | OUT | IN | IN |
| S204E | TRUE | OUT | OUT | OUT | OUT | IN | OUT | IN | IN |
| S204F | FALSE | OUT | OUT | OUT | OUT | IN | IN | IN | OUT |
| S204H | FALSE | OUT | OUT | OUT | OUT | IN | OUT | IN | IN |
| S204T | FALSE | IN | OUT | OUT | OUT | IN | OUT | IN | IN |
| R205D | FALSE | OUT | OUT | IN | OUT | OUT | OUT | OUT | OUT |
| R205F | FALSE | OUT | OUT | IN | OUT | OUT | OUT | OUT | OUT |
| T206A | FALSE | IN | IN | IN | IN | IN | IN | IN | OUT |
| T206C | FALSE | OUT | OUT | IN | IN | OUT | OUT | IN | IN |
| T206K | TRUE | IN | IN | IN | IN | IN | IN | IN | IN |
| T206N | TRUE | IN | IN | OUT | IN | IN | OUT | IN | IN |
| T206R | FALSE | IN | IN | IN | IN | IN | IN | IN | OUT |
| S207T | TRUE | OUT | OUT | OUT | OUT | OUT | IN | OUT | OUT |
| N210E | FALSE | IN | OUT | OUT | OUT | IN | IN | IN | IN |
| N210L | FALSE | OUT | OUT | OUT | OUT | OUT | IN | OUT | OUT |
| A212T | FALSE | IN | OUT | OUT | OUT | IN | IN | IN | IN |
| D213N | FALSE | OUT | OUT | OUT | IN | OUT | OUT | OUT | OUT |
| A217C | FALSE | OUT | OUT | OUT | OUT | OUT | OUT | IN | IN |
| A217N | FALSE | IN | IN | OUT | OUT | IN | OUT | IN | IN |
| A217Y | TRUE | IN | OUT | OUT | OUT | IN | OUT | OUT | OUT |
| E218Q | FALSE | OUT | OUT | OUT | OUT | OUT | OUT | OUT | IN |
| T221D | FALSE | OUT | IN | OUT | OUT | OUT | IN | OUT | IN |
| T221L | FALSE | IN | IN | OUT | OUT | IN | OUT | OUT | IN |
| T221Q | TRUE | OUT | IN | OUT | OUT | IN | IN | OUT | IN |
| T221S | TRUE | OUT | IN | OUT | OUT | IN | OUT | OUT | OUT |
| S222G | FALSE | OUT | OUT | IN | IN | OUT | OUT | OUT | OUT |
| S222N | FALSE | OUT | OUT | OUT | IN | OUT | OUT | OUT | OUT |
| S222P | FALSE | OUT | OUT | IN | IN | OUT | OUT | OUT | OUT |
| I223F | FALSE | OUT | OUT | IN | OUT | OUT | OUT | OUT | OUT |
| I223V | FALSE | OUT | OUT | OUT | OUT | OUT | IN | OUT | OUT |
| E224G | FALSE | OUT | IN | OUT | OUT | OUT | OUT | OUT | OUT |
| R225A | FALSE | OUT | OUT | IN | OUT | OUT | OUT | OUT | OUT |
| R225G | FALSE | OUT | OUT | IN | OUT | OUT | OUT | OUT | OUT |
| Q227I | FALSE | IN | OUT | OUT | OUT | OUT | OUT | OUT | OUT |
| Q227L | FALSE | OUT | OUT | IN | OUT | OUT | OUT | OUT | OUT |
| Q227M | TRUE | OUT | OUT | IN | OUT | OUT | IN | OUT | OUT |
| Q228D | FALSE | IN | IN | OUT | IN | IN | IN | IN | IN |
| Q228E | FALSE | IN | IN | OUT | IN | IN | OUT | IN | IN |
| Q228G | FALSE | OUT | OUT | OUT | IN | OUT | OUT | IN | OUT |
| Q228K | TRUE | IN | IN | OUT | OUT | IN | OUT | IN | IN |
| Q228V | FALSE | IN | OUT | OUT | IN | IN | IN | OUT | OUT |
| H229A | TRUE | OUT | IN | OUT | IN | OUT | IN | OUT | OUT |
| H229G | FALSE | OUT | OUT | OUT | IN | OUT | OUT | OUT | IN |

| Variant ID | Exp top 10% | LASSO all | MLP all | LASSO DynaSig | MLP DynaSig | LASSO static | MLP static | LASSO ASA ΔΔG | MLP ASA ΔΔG |
|---|---|---|---|---|---|---|---|---|---|
| H229I | FALSE | OUT | IN | OUT | OUT | OUT | OUT | OUT | IN |
| H229L | FALSE | OUT | OUT | IN | OUT | OUT | OUT | OUT | OUT |
| H229P | FALSE | OUT | OUT | OUT | IN | OUT | OUT | OUT | OUT |
| H229Q | FALSE | OUT | OUT | OUT | OUT | OUT | OUT | IN | OUT |
| H229V | FALSE | OUT | OUT | OUT | OUT | OUT | IN | OUT | OUT |
| P231H | FALSE | OUT | IN | OUT | OUT | OUT | OUT | OUT | OUT |
| P231T | FALSE | OUT | IN | OUT | OUT | OUT | IN | OUT | OUT |
| Q234C | FALSE | OUT | OUT | OUT | OUT | OUT | OUT | IN | OUT |
| Q234E | FALSE | IN | OUT | OUT | OUT | IN | OUT | IN | OUT |
| Q234N | FALSE | IN | OUT | OUT | OUT | IN | OUT | IN | OUT |
| Q234R | FALSE | IN | OUT | OUT | OUT | IN | IN | IN | IN |
| Q234S | FALSE | IN | OUT | OUT | OUT | IN | IN | IN | OUT |
| F235W | FALSE | OUT | OUT | IN | OUT | OUT | OUT | OUT | OUT |
| L242D | TRUE | IN | OUT | IN | IN | OUT | OUT | OUT | OUT |
| L242F | FALSE | OUT | OUT | IN | OUT | OUT | OUT | OUT | OUT |
| L242K | TRUE | OUT | OUT | IN | IN | OUT | OUT | OUT | OUT |
| L242V | TRUE | IN | IN | OUT | OUT | OUT | IN | OUT | IN |
| L246D | FALSE | IN | IN | IN | IN | OUT | OUT | OUT | OUT |
| L246G | FALSE | OUT | OUT | IN | IN | OUT | OUT | OUT | OUT |
| L246Q | FALSE | IN | IN | IN | IN | OUT | OUT | OUT | OUT |
| L246S | TRUE | OUT | OUT | IN | OUT | OUT | OUT | OUT | OUT |
| L246T | TRUE | IN | OUT | IN | OUT | OUT | OUT | OUT | OUT |
| D247L | FALSE | OUT | OUT | OUT | OUT | OUT | OUT | IN | IN |
| D247M | FALSE | OUT | OUT | OUT | OUT | OUT | OUT | IN | OUT |
| D247R | FALSE | OUT | OUT | OUT | OUT | OUT | OUT | IN | IN |
| D247Y | FALSE | OUT | OUT | OUT | OUT | OUT | OUT | IN | IN |
| L248Y | FALSE | OUT | OUT | IN | OUT | OUT | OUT | OUT | OUT |
| K250E | FALSE | OUT | IN | OUT | IN | IN | IN | OUT | OUT |
| K250F | FALSE | OUT | OUT | IN | OUT | IN | OUT | OUT | OUT |
| K250N | FALSE | IN | IN | OUT | IN | IN | IN | OUT | IN |
| T253I | FALSE | OUT | IN | OUT | OUT | OUT | OUT | OUT | OUT |
| N254D | FALSE | OUT | OUT | OUT | IN | OUT | OUT | OUT | OUT |
| N254G | TRUE | OUT | OUT | IN | OUT | OUT | OUT | OUT | OUT |
| N254I | FALSE | OUT | OUT | IN | IN | OUT | OUT | OUT | IN |
| N254P | FALSE | OUT | OUT | IN | OUT | OUT | OUT | OUT | OUT |
| K257G | FALSE | OUT | OUT | IN | OUT | OUT | OUT | OUT | OUT |
| K257N | FALSE | OUT | OUT | IN | IN | IN | IN | OUT | OUT |
| K257Q | TRUE | IN | OUT | IN | IN | IN | IN | OUT | OUT |
| K257T | FALSE | OUT | OUT | OUT | OUT | IN | OUT | OUT | OUT |
| K257V | FALSE | IN | OUT | IN | OUT | IN | IN | OUT | OUT |
| A258F | FALSE | OUT | OUT | OUT | OUT | IN | OUT | OUT | OUT |
| A258V | FALSE | IN | IN | OUT | OUT | IN | IN | OUT | OUT |
| H259I | FALSE | OUT | IN | OUT | OUT | OUT | OUT | OUT | OUT |
| H259S | FALSE | IN | OUT | IN | IN | OUT | OUT | OUT | OUT |
| T260D | FALSE | OUT | OUT | OUT | IN | OUT | IN | OUT | OUT |
| T260K | FALSE | OUT | IN | IN | IN | IN | OUT | OUT | IN |
| T260M | FALSE | IN | OUT | IN | IN | IN | IN | OUT | OUT |
| T260R | FALSE | IN | IN | IN | OUT | IN | OUT | OUT | IN |

Supplementary Table 4 (continued)

| Supplementary Table 4 (continued) | | | | | | | | | |
|---|---|---|---|---|---|---|---|---|---|
| Variant ID | Exp top 10% | LASSO all | MLP all | LASSO DynaSig | MLP DynaSig | LASSO static | MLP static | LASSO ASA ΔΔG | MLP ASA ΔΔG |
| N261H | FALSE | IN | IN | OUT | OUT | IN | OUT | IN | IN |
| R262K | TRUE | IN | OUT | OUT | IN | IN | IN | IN | OUT |
